# Selection in a growing bacterial/yeast colony biases results of mutation accumulation experiments

**DOI:** 10.1101/2021.04.12.439444

**Authors:** Anjali Mahilkar, Sharvari Kemkar, Supreet Saini

## Abstract

Mutations provide the raw material for natural selection to act. Therefore, understanding the variety and relative frequency of different type of mutations is critical to understanding the nature of genetic diversity in a population. Mutation accumulation (MA) experiments have been used in this context to estimate parameters defining mutation rates, distribution of fitness effects (DFE), and spectrum of mutations. MA experiments performed with organisms such as *Drosophila* have an effective population size of one. However, in MA experiments with bacteria and yeast, a single founder is allowed to grow to a size of a colony (~10^8^). The effective population size in these experiments is of the order of 10. In this scenario, while it is assumed that natural selection plays a minimal role in dictating the dynamics of colony growth and therefore, the MA experiment; this effect has not been tested explicitly. In this work, we simulate colony growth and perform an MA experiment, and demonstrate that selection ensures that, in an MA experiment, fraction of all mutations that are beneficial is over represented by a factor greater than two. The DFE of beneficial and deleterious mutations are accurately captured in an MA experiment. We show that the effect of selection in a growing colony varies non-monotonically and that, in the face of natural selection dictating an MA experiment, estimates of mutation rate of an organism is not trivial. We perform experiments with 160 MA lines of *E. coli*, and demonstrate that rate of change of mean fitness is a non-monotonic function of the colony size, and that selection acts differently in different sectors of a growing colony. Overall, we demonstrate that the results of MA experiments need to be revisited taking into account the action of selection in a growing colony.

## Introduction

Mutations create genetic diversity in a population. The diversity and the relative frequency of beneficial, neutral, and deleterious mutations in a given environment dictate the ability of a population to respond to an adaptive challenge. The most direct way of studying this diversity is with the help of a mutation accumulation (MA) experiment. In these experiments, a population is allowed to expand to a fixed size, and forced to go through a severe bottleneck. In this bottleneck, for instance, only one, randomly chosen individual (or one individual from each sex) is allowed to progress to the next generation. The mutations so accumulated are believed to be largely independent of selection, and hence, represent the true spectrum of mutations in an organism.

Previously, mutation accumulation experiments have been performed in a large number of species, including multicellular eukaryotic species like *Arabidopsis thaliana* (Ossowski et al., 2010, Rutter et al., 2012), *Caenorhabditis elegans* (Baer et al., 2005, Denver et al., 2012), *Drosophila melanogaster* (Haag-Liautard et al., 2007, Keightley et al., 2009, Schrider et al., 2013, Mukai, 1964, Mukai et al., 1972). MA experiments have also been performed on single-celled eukaryote *Saccharomyces cerevisiae* (Lynch et al., 2008, Dickinson, 2008) and *Chlamydomonas reinhardtii* (Ness et al., 2012) and with bacteria such as *Escherichia coli* (Kibota & Lynch, 1996) and *Salmonella typhimurium* (Andersson & Hughes, 1996). One of the most consistent patterns from MA experiments has been that as the number of transfers progress, the mean fitness of independent lines decreases, and the variance of the mean fitness between independent lines increases (Kibota & Lynch, 1996).

These experiments have provided us with key insights into mutational events of an organism. These insights include (a) the overall mutation rate (Caballero & Keightley, 1998, Kibota & Lynch, 1996), (b) the general mutational pattern (Kraemer et al., 2017, Estes et al., 2004, Dillon et al., 2018, Ann-Marie Waldvogel, 2020), (c) estimating the shape of the distribution of fitness effects of mutations (Bondel et al., 2019, Bosshard et al., 2017, Mrudula Sane, 2020). While in eukaryotes, the longer generation times and larger body size permit separation between every generation; the same is not possible in prokaryotes like bacteria, or single-celled eukaryotes like yeast. As a result, mutation accumulation experiments in yeast and bacteria are performed by growing cells for several generations on solid agar. After a period of growth (~20-25 generations), a colony is selected at random and cells from it spread on agar plates (**Figure 1**). This process is repeated for a number of transfers (typically *~*100). Since selection of an individual which forms the colony to be picked is completely randomized (by picking a random colony), the effects of selection are thought to be minimized in such an experiment. One of the reasons attributed to this is that since most cell divisions (and consequently mutations) appear in the last generation, a bottleneck is imposed before selection can act on these mutations (Lee et al., 2012). The repeated bottlenecks then drive accumulation of mutations primarily via drift (Katju & Bergthorsson, 2019). However, to what extent does action of selection dictate the growth phase of a colony is not known (Heilbron et al., 2014).

**Figure 1.**
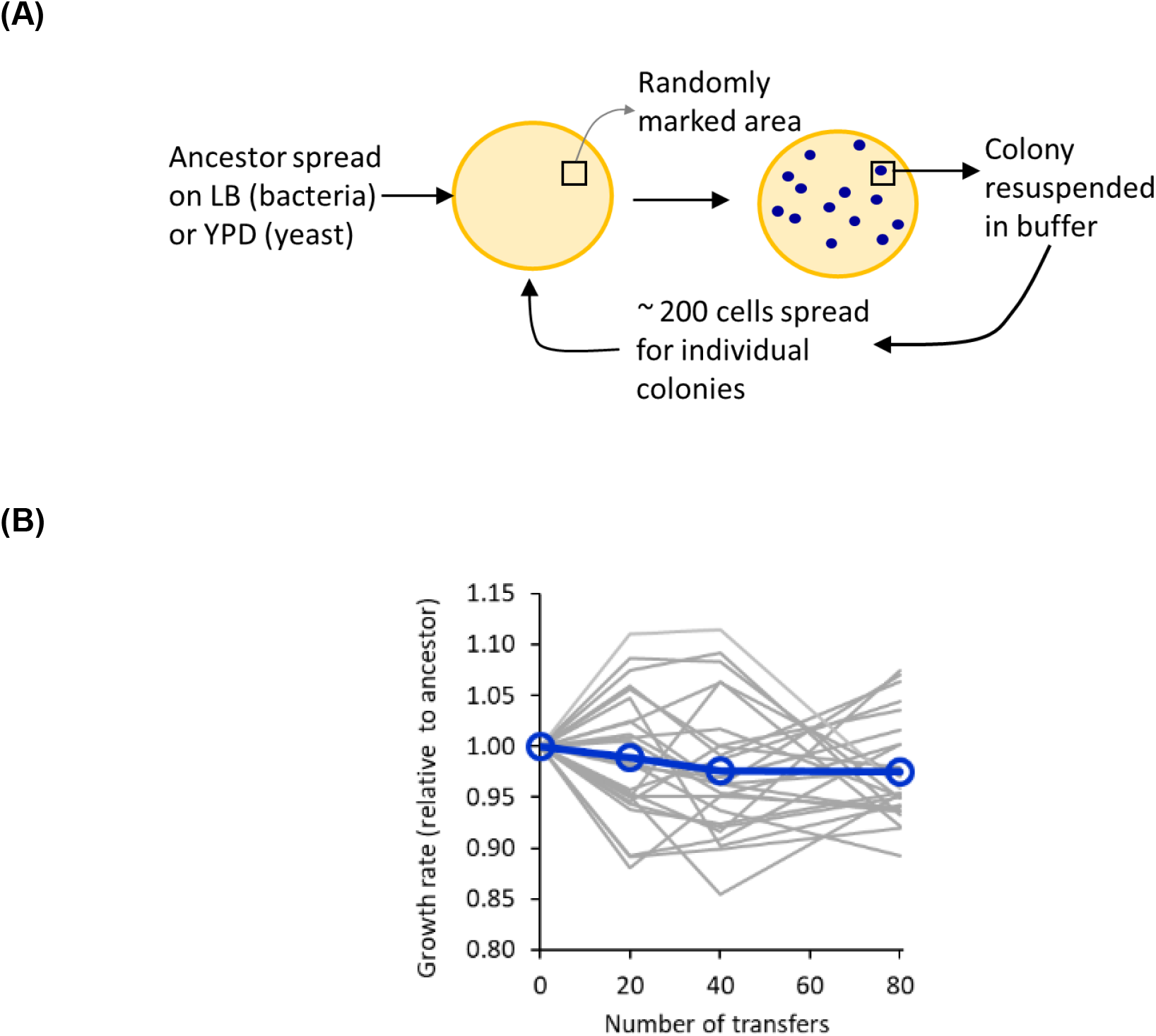
**(A) MA experiment methodology for yeast and bacteria.** An area of the plate is marked and *N* cells spread on the plate. A colony in the marked area is picked and the process repeated. **(B) Change in growth rate in a mutation accumulation experiment.** Twenty-two lines of *E. coli* were propagated in an MA experiment. Growth rate of the lines at regular intervals was recorded (grey lines) and is represented as relative to that of the respective ancestor. The mean of the 22 lines is represented as dark blue curve. All measurements are averages of three repeats. The standard deviation in the growth rate measurement is less than 0.03%.

In this work, we show that selection plays a significant role in dictating the evolutionary dynamics in an MA experiments with bacteria and yeast. The observed mutational spectrum from the MA experiments does not mimic the underlying distribution from which mutations occur randomly. In fact, as a manifestation of an MA experiment, deleterious mutations are significantly under-represented, and beneficial mutations greatly over-represented. Interestingly, there is no significant change in the distribution of fitness effects (DFE) as observed in an MA experiment, and the underlying theoretical distribution. Parameters such as mutation rate, estimated from an MA experiment, are also influenced by selection. We demonstrate that inferences about mutational spectrums of bacteria/yeast from mutation experiments are influenced by selection.

## Methods

### Simulations

#### Simulation of colony growth

The growth of a colony is simulated starting from a single individual. The growth of a colony is divided into two phases. In the first, starting from the founder individual, growth is modelled stochastically upto a size of 60,000 individuals. This phase is followed by a deterministic phase of growth, upto the final colony size.

The stochastic phase is simulated as follows. The cell division time of the founder of the colony is assumed to be 100 minutes. Although members of an isogenic bacterial/yeast populations exhibit a distribution of division times, we assume that all isogenic cells divide at the same time.

When cell division occurs, a progeny identical to the parent is generated with a probability (1-μ). The progeny has a mutation with probability μ. The mutation is beneficial with probability *b*, and deleterious with probability (1-*b*). Neutral mutations are not taken into account in our model. It is however known that neutral mutations can influence the evolutionary trajectories of population. An exponential distribution is assumed for both beneficial and deleterious mutations. The distributions are represented by parameters λ_b_ and λ_d_ for beneficial and deleterious mutations, respectively. It has, however, been proposed that distributions describing deleterious mutations are more complex (Brajesh et al., 2019, Elena et al., 1998, Neher, 2013). The parameters for the simulations are given in **Table 1**.

**Table 1.** List of parameters and their values in the simulation.

| Parameter | Value |
| --- | --- |
| Mutation rate, $\mu$ | 0.001 per cell per division |
| Fraction of mutations which are beneficial, $\alpha$ | 0.25 |
| Exponential distribution parameter for beneficial mutations, $\lambda_b$ | 1 |
| Exponential distribution parameter for beneficial mutations, $\lambda_d$ | 1 |
| Division time of the ancestor, $1/f_0$ | 100 minutes |

Once an individual is simulated to have acquired a beneficial mutation, the magnitude of the beneficial mutation is identified by a random draw from the exponential distribution with parameter λ_b_. The same strategy is identified for deleterious mutations. After acquisition of a mutation, the division time of the progeny is updated accordingly.

In this phase of growth, the individual closest to its time of division is identified and chosen for division in the manner described in the paragraph above. For all other cells, the time to division is updated accordingly. This process is continued till the population reaches a size of 60,000.

After this initial stochastic growth of the population to a size of 60,000, further expansion to a full colony size is modelled via a deterministic scheme. MA experiments are thought to minimize the effect of selection because the maximum number of cell divisions (and hence mutations) occur in the last generation of growth. As a result, selection has no (little) time to act on this genetic variation. This was simulated by the following strategy in the deterministically modelled phase of growth.

The genotypes in the colony at size 60,000 are recorded and distributed in bins as described in **Table 2**. Further growth is modelled in steps, where each step is one generation time of the founder genotype. In this step, newer mutations in the founder genotype are allowed to arise. Since the number of individuals carrying a beneficial or deleterious mutations is small, we assume that no newer mutations arise in this pool. For example, if the number of founder genotype is *P* at a certain instant. In the next generation, *P*(1-μ) identical individuals are added to the number of individuals of this genotype. *P*μ*b* individuals with beneficial mutations are added to the population; and *P*μ(1-*b*) individuals with deleterious mutations are added to the population. The fitness values of the mutants in this round of growth is allotted as per the exponential distributions defined by λ_b_ and λ_d_. This process is continued till the population size reaches *K.*

**Table 2.** Fitness bins in which population frequency was stored when colony size reached 60,000.

| Deleterious | Beneficial |
| --- | --- |
| $< 0.75 f_0$ | $1 - 1.01 f_0$ |
| $0.75 - 0.80 f_0$ | $1.01 - 1.02 f_0$ |
| $0.80 - 0.85 f_0$ | $1.02 - 1.03 f_0$ |
| $0.85 - 0.90 f_0$ | $1.03 - 1.04 f_0$ |
| $0.90 - 0.95 f_0$ | $1.04 - 1.05 f_0$ |
| $0.95 - 1.00 f_0$ | $> 1.05 f_0$ |

Individuals in each bin, as per Table 2, are assumed to grow with a growth rate equal to the average fitness of the bin. A set of coupled ODEs of the form (1) are solved numerically till the population size reaches *K*, using ode45 on MATLAB.

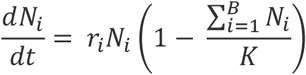

Where, *K* is the maximum colony size. In this work *K* is taken to be 1.1 x 10^8^. As a result, in simulations where propagation of colonies takes place at population size 10^6^, the colony size is only 0.01 times the maximum size of the colony on the particular media.

#### Model assumptions

In addition to the above, the following changes were made to the model and their effect quantified.

It is known that division times of isogenic populations are represented by log-freschet distributions (Pugatch, 2015). In the representation above, all isogenic individuals are assumed to divide at the exact time. This assumption was relaxed when the division time for an individual from an isogenic population was chosen randomly from a normal distribution with a standard deviation of five percent around the mean.

Recent work on microbial systems has demonstrated that epistasis influences the dynamics of adaptation (Gautam Reddy, 2020, Good & Desai, 2015, Kryazhimskiy et al., 2014). However, little is known about the statistics of epistasis at a global level in a cell. We, therefore, do not include the effect of epistasis in our model, and assume additive effect of mutations on fitness.

Effect of neutral mutations. The effect of a mutation on the fitness of an organism can be beneficial, deleterious, or neutral. While neutral mutations do not change the fitness of the individual, they have the potential of changing the adaptive trajectory of the organism (Draghi et al., 2010, Wagner, 2012, Wagner, 2005, Wagner, 2008). In this context, it has been suggested that an intermediate fraction of all mutations being neutral can speed up the adaptive response of a population, and can thus be under secondary selection (Draghi et al., 2010). We do not consider this effect in our model.

#### Simulating MA trajectory

Colony growth, using the above described simulation scheme, was simulated for three colony sizes: 10^6^, 10^7^, and 10^8^. For each size, the colony was simulated to grow to a respective size. At this point, the number of individuals carrying beneficial (N_b_) and deleterious (N_d_) mutations along with their respective fitness was stored. After growth to the appropriate colony size, one individual was picked randomly from the colony. This individual was then considered as the founder for the next transfer in an MA experiment. This process was continued for 2500 transfers. The mean fitness of the colony at each transfer was recorded.

In simulations where the effect of selection was ignored. The acquisition of mutations and the distribution of fitness across members of a colony was recorded. However, it was assumed that these mutations did not change the phenotype of the individual cells, and each cell divided at the same rate as that of the ancestor.

### Experiments

#### Media and reagents

Luria-Bertani (LB) media was used for all experiments

#### Mutation Accumulation Evolution Experiment

*E. coli* K12 MG1655 (ATCC 47076) was revived from freezer stock in LB and grown overnight at 37°C and 250 rpm. MA evolution lines were started using 2ul of overnight grown culture to make spots and streak using sterile toothpicks. A single colony was used as a founder for all mutation accumulation lines. At every transfer, prior to spreading the cells on the plate, an area was marked. The colony in (or closest to) the marked area was chosen for the next transfer for each line. Thirty two independent lines were propagated for each of the following five conditions: (a) transfer after 8 hours of growth on LB plates at 37 deg C. These 32 lines are referred to as “small1” (s1) to “small32” (s32); (b) transfer after 12 hours of growth on LB plates at 37 deg C. These 32 lines are referred to as “medium1” (m1) to “medium32” (m32);; (c) transfer after 24 hours of growth on LB plates at 37 deg C. These 32 lines are referred to as “l1” (l1) to “l32” (l32); (d) transfer after 24 hours of growth at 27 deg C on LB plates, where cells from the outermost edge are propagated further. These 32 lines are referred to as “edge1” (e1) to “edge32” (e32); and (e) transfer after 24 hours of growth on LB plates at 37 deg C, where cells from the center of the colony are propagated further. These 32 lines are referred to as “center1” (c1) to “center32” (c32).

The MA evolution experiment was carried out for a total of 100 transfer for each of the five groups of 32 lines. For the 32 lines (s1-s32), the MA experiment was done for 150 transfers. The evolved lines were stored after every 50 transfers at −80°C in 25% glycerol.

#### Growth rate estimation of MA lines

For calculating the relative fitness of the MA lines, cells from the MA lines and the ancestor were revived from the freezer stock in 2ml LB. The cultures were then grown for 12 hours at 37°C and 250 rpm. The cultures were then sub-cultured 1:100 into fresh 2ml LB media. A volume of 150 μL of these cultures were transferred to a 96-well clear flat-bottom microplate (Costar) in triplicates. The cultures were grown at 37°C in an automatic microplate reader (Tecan Infinite M200 Pro), until they reached stationary phase. OD600 readings were taken every 30 minutes with 10 minutes of orbital shaking at 5mm amplitude before the readings. A gas permeable *Breathe-Easy* (Sigma-Aldrich) sealing membrane was used to seal the 96-well plates. Growth rates were calculated as described in (Mrudula Sane, 2020).

#### Temperature stress experiment

Cells were revived from the freezer stock in 2 ml LB and allowed to grow at 37 deg C for 24 hours. The saturated cultures were then diluted in 2 ml water to an optical density of 0.2. The diluted cultures were then immersed in a hot water bath at temperature 50 deg C for one hour. After the heat shock, the cultures were (a) spot plated at a dilution 1:1000 on LB plates, and (b) sub-cultured 1:1000 in 2 ml LB and allowed to grow at 37 deg C. Optical density at 600 nm of these cultures was recorded after a period of four hours of growth.

## Results

### Increase in fitness of mutation accumulation (MA) experiment lines

We performed MA experiments by streaking *E. coli* (22 lines) on LB plates for a total of 80 transfers. This work started with the observation that most lines of our MA experiment exhibited a growth rate higher than that of the ancestor. Growth rate (proxy for fitness) of the evolved lines at 20, 40, and 80 transfers, is shown in **Figure 1B**. Somewhat surprisingly, even as the mean growth rate of the 22 lines decreased, 10 out of the 22 lines exhibited an increase in the growth rate by 40 transfers. The number of lines, which exhibited fitness greater than that of the ancestor, decreased to seven at transfer 80. Of all data points, which exhibited a statistically significant change in growth rate between two successive points of measurement, ~45% exhibited an increase in growth rate, and ~55% exhibited a decrease. The ratio between the two is not indicative of the ratio of beneficial to deleterious mutations believed to be available to a genotype (Perfeito et al., 2007).

While increase in growth rate in MA experiments has been reported previously (Hall et al., 2013, Dillon & Cooper, 2016, Krasovec et al., 2017, Keightley & Caballero, 1997, Zeyl & DeVisser, 2001, Kuroki et al., 1993), it has been attributed to action of drift (Katju & Bergthorsson, 2019). Repeated bottlenecks in the population, after the period of colony growth (~25 generations) are believed to negate the effect of selection. However, this idea remains based largely on qualitative intuition. This result formed the basis for our simulation and experiments, as described in the rest of the manuscript.

### Model description to mimic colony growth

Starting with an ancestor with fitness (1/t_o_) (where *t_o_* is the division time), growth of a colony is mimicked (**Figure 2A**). Each progeny inherits the ancestor’s genotype (and fitness) with a probability (1-*μ*), where *μ* is the mutation rate per cell per division. In case one or both the progenies acquires a mutation, the mutant progeny is allotted a fitness as per the following logic. Of all mutations occurring, a fraction *b* are assumed to be beneficial. As a result, a fraction (1-*b*) are assumed to be deleterious. Neutral mutations are not considered in our model. The DFE of beneficial and deleterious mutations is assumed to be exponential in nature (Brajesh et al., 2019, Wrenbeck et al., 2017, Orr, 2010), and is represented by *λ_b_* and *λ_d_*, respectively. Depending on the nature of mutation (beneficial or deleterious) acquired by the mutant, a random number from the corresponding exponential distribution is drawn and assigned to be the progeny’s fitness. A progeny with fitness *f_mut_* undergoes division 1/*f_mut_* time post-birth. When the colony consists of *i* number of cells, the individual cell closest to its division time is chosen for division. The time to division of all other cells in the colony is adjusted accordingly. This phase of growth is modelled till the colony size reaches 60,000 individuals.

**Figure 2.**
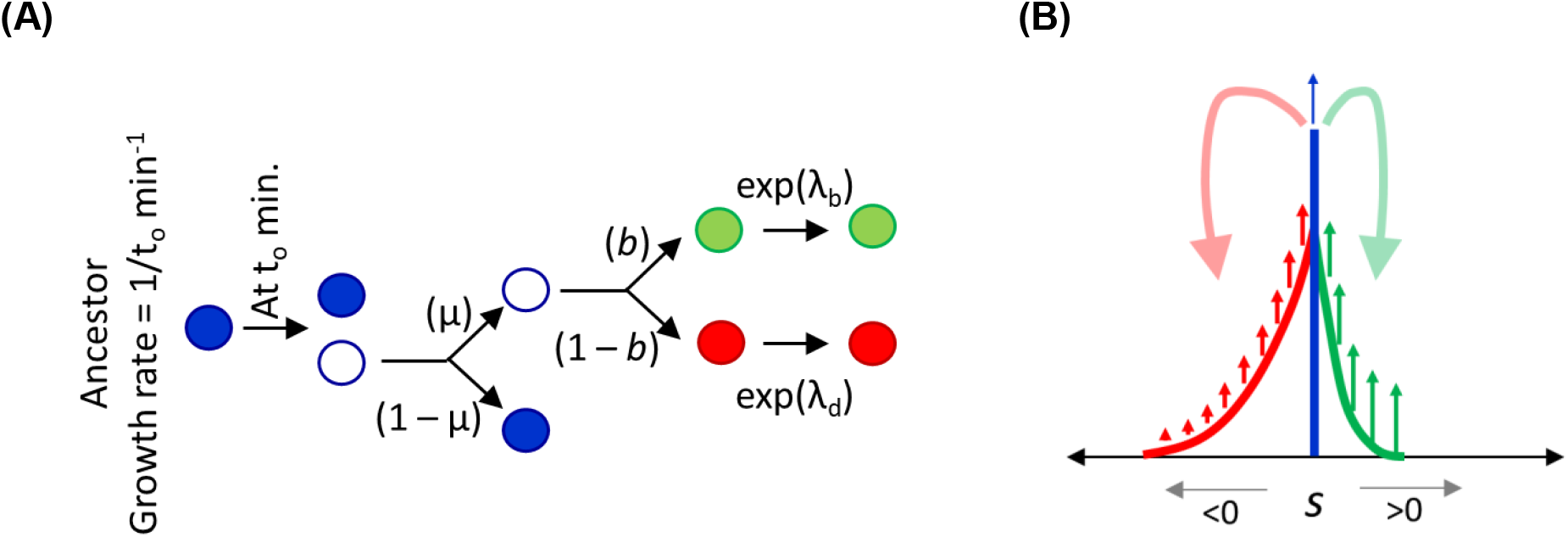
**(A)** Cell division model. At time to, a cell (with growth rate 1/to) was assumed to divide into two. The acquired mutation is beneficial with probability *b*. Neutral ismutations were not considered in the model. The selection coefficient of a mutation was drawn randomly from an exponential distribution with parameters λb and λd, for beneficial and deleterious mutations, respectively. **(B)** Colony growth model. After growth to a size of 60,000 individuals as described in (A), the growth of individuals of genotype was modeled deterministically. Red (green) indicates the individuals with deleterious (beneficial) mutations. Blue indicates individuals with no mutations. As the colony size grows, in each generation, newer mutations occur in the parent genotype, and this contributes to the pool of beneficial and deleterious mutants. The number of such mutants is computed simply as *Nμ*, where *N* is the number of individuals of ancestral genotype in that generation. The fitness effects of such mutants were allotted as per the exponential distribution describing beneficial and deleterious mutations.

At this time, the various genotypes present in the colony are assumed to grow deterministically till the population reaches the final size of the colony *N*_f_ (see methods for more details). As shown in **Figure 2B**, the individuals with deleterious mutations grow at a rate lower than the ancestor, while those with beneficial mutations grow at a rate higher than that of the ancestor. Mutations in the ancestor contribute to the pool of beneficial and deleterious mutants. For example, in one generation if *N* ancestor cells divide, *N*μ*b* beneficial mutants and *N*μ(1-*b*) deleterious mutants are given birth. The fitness of these newly produced mutants is allotted deterministically as per the DFE. Since the number of mutant genotypes in a growing colony is small, the number of mutations arising in these individuals is small, and hence, not considered in the model.

In this representation, we make several assumptions. First, cell division times are known to be distributed around a mean (Wang et al., 2010, Pugatch, 2015), and not be a constant for an isogenic population. Second, exponential distribution parameters, *λ_b_* and *λ_d_*, for beneficial and deleterious mutations, will be a function of the strain, fitness, and the environment in which the experiment is performed (Brajesh et al., 2019). However, we consider these parameters to be constant for the duration of the simulation. Moreover, previous work has also suggested that the DFE for deleterious mutations have shapes other than an exponential distribution (Brajesh et al., 2019, Neher, 2013).

### Selection in action during growth in a colony

Mutation accumulation experiments assume that the action of selection is minimal due to the repeated bottleneck in the experimental setup every ~20-25 generations. A typical colony grows from a size *N* = 1, to *N* = 10^8^, and in this period several beneficial and deleterious mutations arise. In fact, several genotypes compete in a growing colony (**Figure 3A**). This distribution is consistent with the variation observed experimentally in rich media conditions (**Figure 3B**).

**Figure 3.**
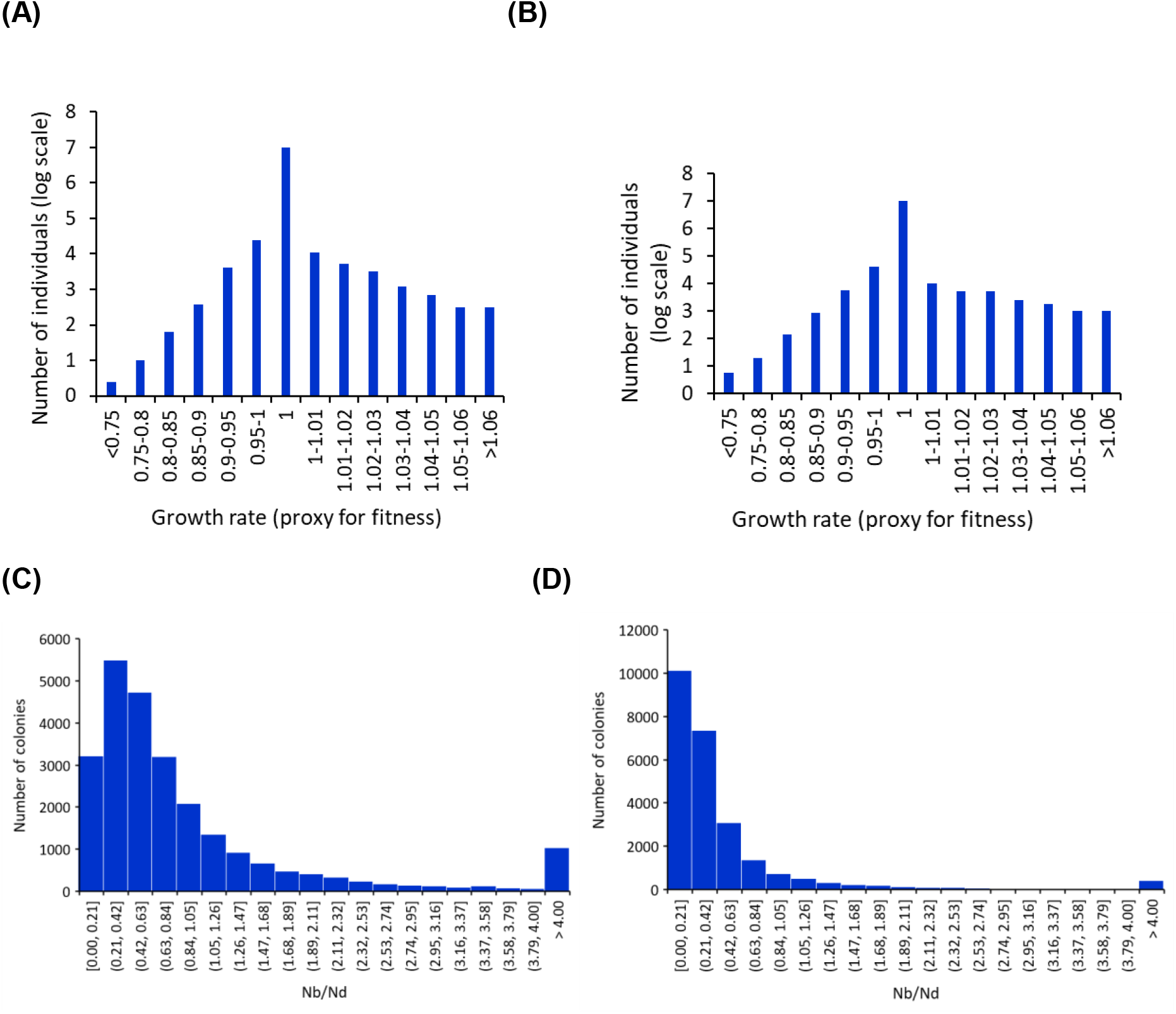
**(A)** Genotypes with different phenotypes compete in a growing colony. Distribution of fitness phenotype in a single representative colony. The fitness of the ancestor is normalized to one. All individuals with fitness >1 (<1) carry beneficial (deleterious) mutation(s). **(B)** The distribution of mutations remains qualitatively the same when the cell division time is not exact but a normal distribution around the mean value. The division time was assumed to be a normal distribution with a standard deviation of 5% of the mean. Data for a single colony. **(C)** Distribution of the number of individuals with beneficial and deleterious mutants in a colony. The simulation in (A) was repeated 25000 times, and the frequency distribution of the ratio of individuals with beneficial to deleterious mutations plotted. **(D)** Frequency distribution of the ratio of number of individuals with beneficial and deleterious mutations in a colony, if all mutations were neutral.

However, the process of mutations is inherently stochastic. Effects like the timing and nature of mutations deeply impact the final genetic structure of the colony. Thus, the number of beneficial and deleterious mutants present in a colony vary significantly from one colony to another (**Figure 3C**). The distribution of the ratio N_b_/N_d_ (ratio of the number of individuals with a beneficial mutation and the number of individuals with a deleterious mutations) across a simulation of 25000 colonies exhibits a wide distribution. This is significantly different from the distribution of Nb/Nd expected if beneficial mutations did not have any selection effect in a colony environment (**Figure 3D**). As a result of the long-tail, the mean of the distribution of Nb/Nd in Figure 2B is approximately 1.4. The null expectation of this ratio, in the absence of selection is 0.33. This bias is largely introduced by the colonies in which beneficial mutations appear relatively early in the colony growth. Thus, selection acting in a growing colony qualitatively changes the distribution of the number of individuals carrying a beneficial vs deleterious mutations.

### Mimicking an MA experiment

From the simulation of a colony, we extend our analysis to mimic an MA experiment. For this purpose, we follow the following strategy. Starting from an ancestor, we simulate colony growth and have estimates for N_b_, N_d_, and the distribution of fitness effects. From this distribution, we pick an individual randomly, and similarly simulate the kinetics of colony growth again. Selection acts in a growing colony with various genotypes competing against each other. The likelihood of picking a particular fitness phenotype as a founder is simply proportional to the frequency of the genotype with the corresponding phenotype. By repeating this process, we mimic an MA experiment.

The mean fitness of an MA line decreases linearly with time (**Figure 4A**). The changes in fitness are consistent with previous experimental observations (Kibota & Lynch, 1996). A large fraction of MA lines exhibit an increase in fitness, in the early parts of the experiment. The fraction of MA lines with increased fitness exhibits a characteristic pattern, and the variance of fitness among MA lines increases with the number of transfers (**Figure 4B and 4C**). These trajectories are significantly different if selection were not acting in a growing colony (**Figure 4D-F**).

**Figure 4.**
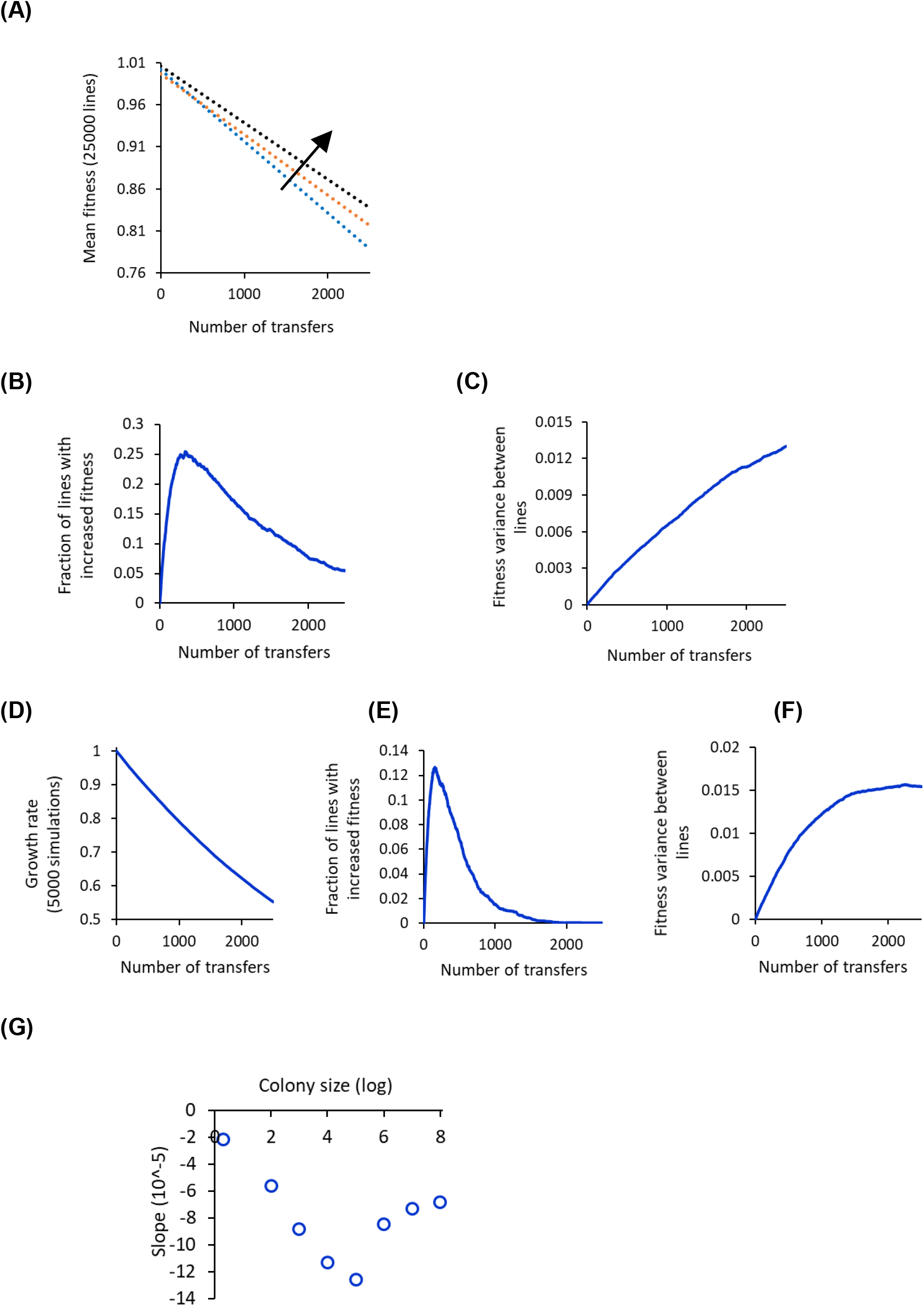
**(A)** Mean fitness of a colony decreases with increasing transfers in an MA experiment. An average profile of 25000 colonies shows a linear decrease in fitness with increasing number of transfers. Blue curve scenario when propagation from one plate to another was done when colony size was 10^6^, orange for 10^7^, and black for 10^8^. The arrow indicates the direction of increasing colony size. **(B) The fraction of MA lines which show fitness greater than that of the ancestor as a function of transfer number.** Stochastic effects and selection lead to ~25% of the lines with fitness greater than that of the ancestor. With increasing number of transfers, this fraction decreases to about five percent. **(C) The variance of fitness across lines increases with increasing number of transfers.** Slope at different colony sizes in an MA experiment shows a non-monotonic trend. Average of 25000 independent MA lines. **(D-F)** Plots of 4A, B, and C assuming that selection was not affecting the growth phenotype in the growing colony. (D) mean growth rate of the colony, (E) fraction of lines with fitness greater than that of the ancestor, and (F) variance between fitness values of the individual lines is shown as a function of number of transfers. **(G) Slope of fitness vs. number of transfers in an MA experiment with different colony sizes.** The slot exhibits a non-monotonic behaviour with the slope minimum for intermediate colony sizes.

MA experiments are thought to be independent of bias since most of the mutations are thought to take place in the last generation of colony growth. As a result, selection does not get a chance to act on these mutations. However, colonies where mutations occur early are not free from the action of selection, and they significantly alter the growth kinetics of beneficial and deleterious mutants.

Two factors dictate the relative strengths of selection and drift acting on a colony. As the colony grows to a greater size, selection has greater time to act on the existing genotypes in the colony (particularly those mutations that arose in the first few generations). On the other hand, greater colony size implies that a large number of divisions (and hence mutations) take place in the last generation – and hence, selection cannot act on it.

The trajectory of mean fitness across 25000 independent MA lines for different colony size at the time of transfers demonstrates that selection is stronger in a large colony, compared to a smaller one (**Figure 4A**). In fact, our simulations predict that the slope of the mean fitness trajectory (across 25000 MA lines) exhibits a non-monotonic profile (**Figure 4G**). If cells are transferred from one plate to another when the colony size is small, selection has little time to impact the relative frequency of the different genotypes in the colony. In this phase of the colony growth, competition for resources is limited, and the colony growth exhibits an exponential growth. However, as the colony increases in size, competition for resources intensifies, and growth is linear in nature. In such a setting, selection becomes far more important. As a result of the two phasic growth of a colony CFU, the relative intensity and effects of selection and drift change with colony growth.

### DFE from an MA experiment exhibits bias towards beneficial mutations

We now return to our original question of how does selection in an MA experiment bias derivation of parameters associated with mutational spectrum of an organism. In particular, we are interested in investigating the (a) DFE and (b) mutation rate estimates from an MA experiment data. We first discuss (a).

DFE of beneficial and deleterious mutations are recorded from the simulations via two parameters. First, of all the mutations occurring (seen in an MA simulation), what fraction are beneficial. Second, what are the corresponding DFEs for beneficial and deleterious mutations. For both these parameters, we extract observations from the MA simulations and compare with the underlying parameters/distributions, which formed the basis of our MA simulations.

The action of selection biases MA experiments towards a greater observation of beneficial mutations. From an underlying distribution of 25% mutations being beneficial, action of selection in an MA experiment simulation changes that to 57%. This bias perhaps also explains the rather surprisingly high fraction of mutations being beneficial, as reported recently (Mrudula Sane, 2020).

Estimate of the DFE for beneficial and deleterious mutations were done by fitting a *λ* parameter on the observed distribution of first mutations in simulated MA experiments. The fitted parameters were then compared with the underlying theoretical distribution (λ in **Table 1**). For both, beneficial and deleterious mutations, the observed distributions from the MA experiment mimic the underlying distributions from which mutations are drawn (**Figures 5A** and **5B**).

**Figure 5.**
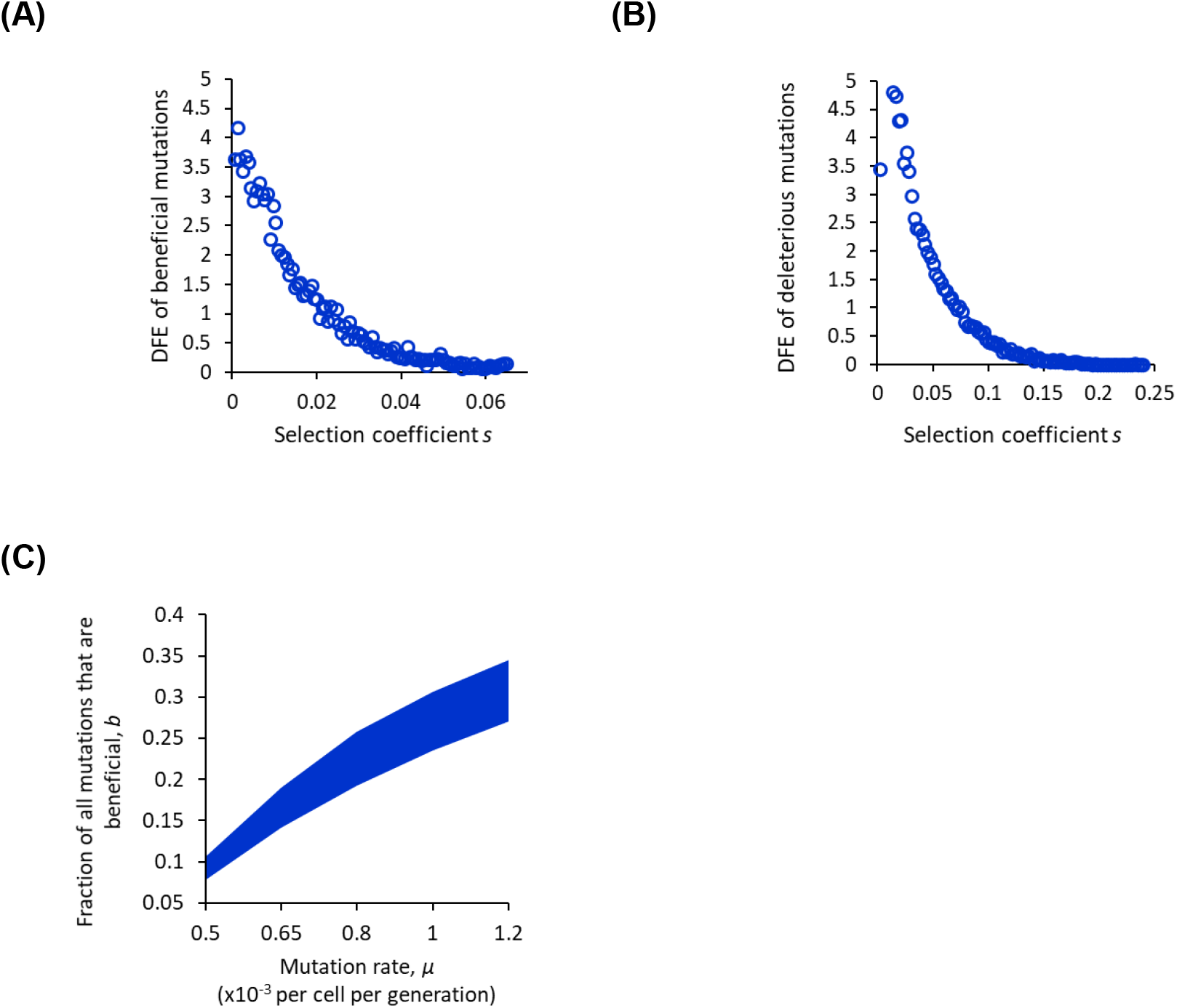
MA experiments accurately predict the DFE of **(A)** beneficial and **(B)** deleterious mutations. Distribution of 25000 beneficial mutations acquired and propagated in ancestor. The observed distributions can be represented as an exponential distributions with λ equal to 67.1 (A) and 26.97 (B) and are statistically identical with the *λ* values in Table S1 (p-value <0.01). **(C)** A large section of the two-dimensional parameter space explains the MA simulations. The highlighted region indicates parameter values (*μ, b*) which explain the MA simulation trajectory in Figure 3A for colony size 10^7^ with 95% confidence.

Mutation rates of species have been predicted from MA experiments (Andersson & Hughes, 1996, Kibota & Lynch, 1996). However, these studies assume that beneficial mutations are sufficiently rare that they are not picked up due to selection in MA experiments. In addition, mutations with large deleterious affects (lethal mutations) can be ignored from the analysis. Thus, the assumption that it is deleterious mutations, which are identified in MA experiments. However, our simulation results show this to be not true.

Therefore, a set of parameters define the outcome of an MA experiment. These parameters include (a) mutation rate, *μ*, (b) fraction of mutations which are beneficial, *b*, (c) parameter *λ_b_* describing distribution of beneficial mutations, and (d) parameter *λ_d_* describing distribution of deleterious mutations. This parameter space constitutes a 4-dimensional space, and the challenge is to identify regions of this space, which describes the observations from an MA experiment. We fix *λ_b_* and *λ_d_*, and scan the two-dimensional space described by mutation rate *μ*, and fraction of mutations that are beneficial, *b* to fit the data from the MA experiment simulation. As shown in **Figure 5C**, the estimate of mutation rate *μ* from the MA experiment simulation is dependent on the parameter *b*. With increasing *b*, higher mutation rates are able to explain the observed MA line trajectory, as observed in an experiment/simulation. Since *b* is largely unknown for a particular environment, estimates of *μ* from an MA experiment are quite challenging.

### Testing model predictions in an MA experiment

An MA experiment was performed with 160 parallel lines of *E. coli*. The lines were separated into five categories, with 32-identically treated lines in each. The first three categories were MA lines propagated at different colony sizes. These lines are called small (lines s1-s32), medium (m1-m32), and large (l1-l32). The other two categories were large colonies where the cells propagated from one transfer to the next were picked from edge of the colony (lines e1-e32) and centre of the colony (lines c1-c32).

The distribution of fitness of the 32 lines at transfers 50, 100, and 150 is shown in **Figure 6**. The mean fitness of these lines decreases as the number of propagations increase (**Figure 7A)**.

**Figure 6.**
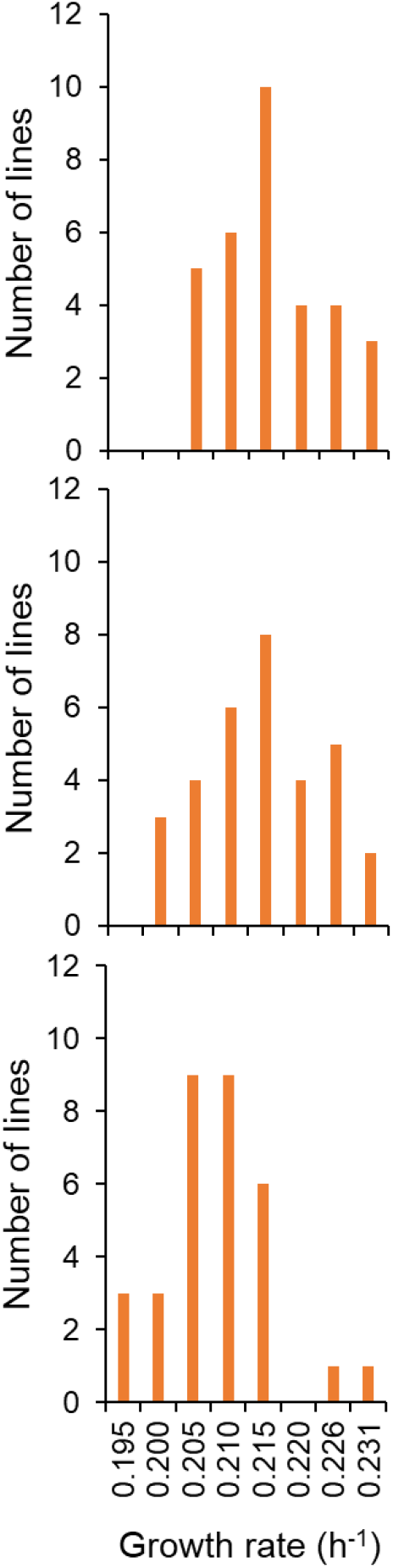
Distribution of fitness among the 32 small lines (s1-s32). Top panel represents the distribution after 50 transfers, the middle panel after a 100 transfers, and the lower panel after 150 transfers.

**Figure 7.**
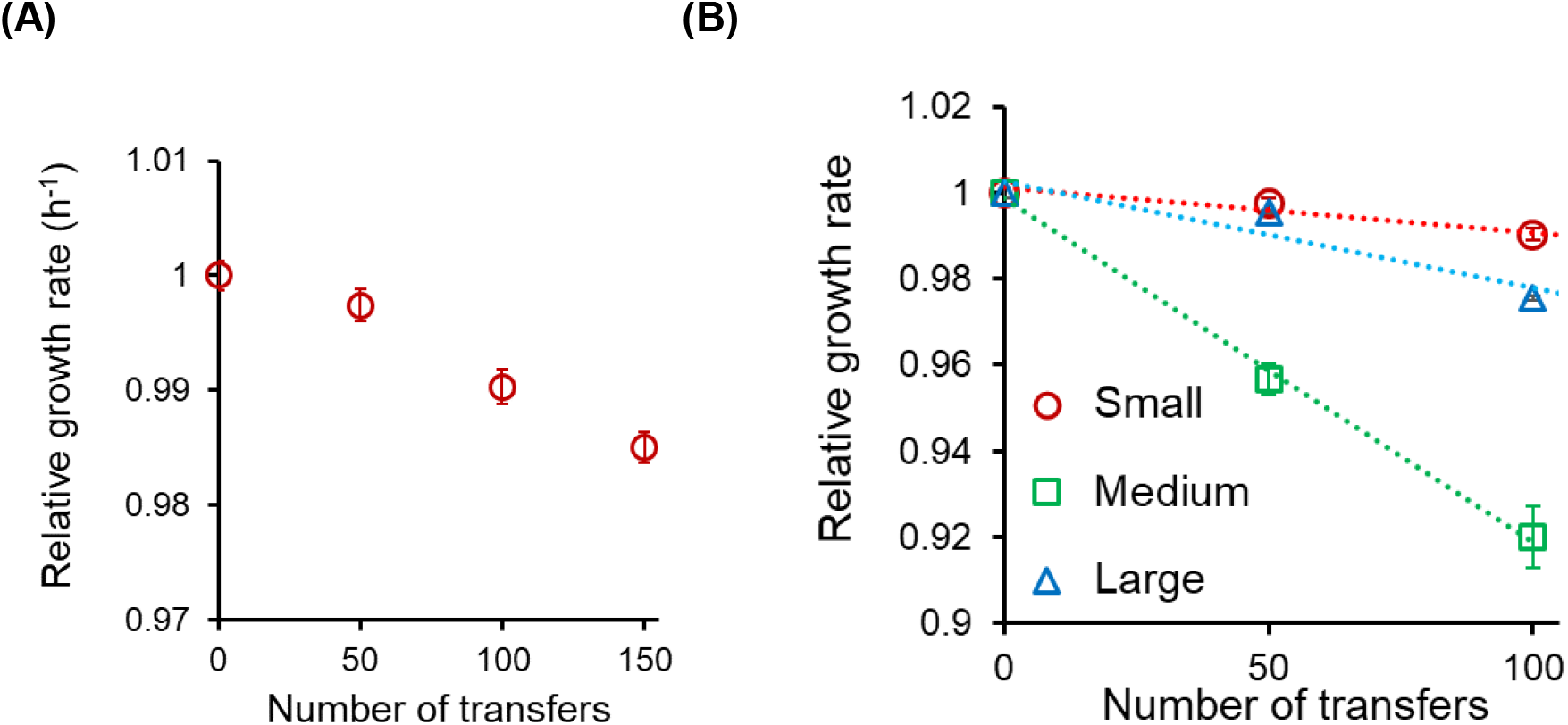
**(A)** The mean fitness of the 32 small lines (s1-s32) decreases linearly with the number of transfers. **(B)** Mean fitness of the 32 MA lines decreases non-monotonically as a function of the colony size. The rate of decrease in the largest (in magnitude) for medium colony size (m1-m32) (green squares), compared to small (s1-s32) (red circles), and large (l1-l32) (blue triangle).

Consistent with our model predictions, the rate of decrease of the mean fitness is the greatest in the lines of medium size, compared to that in the large and small colony size propagations. (**Figure 7B**). Although the linear behaviour of the decrease in fitness has been previously reported in MA experiments with bacteria, the non-monotonic nature of this decrease has not been reported before. Our model shows that this non-monotonic behaviour is because of the change in the relative strength of selection and drift, as colony size changes.

We explicitly test the assumption that different sectors of a growing colony are under different selection pressures. The microenvironments of a growing colony are known to be quite distinct from each other, and hence, the individuals in these sectors are likely under different selection pressures (Cap et al., 2010, Cap et al., 2009). For example, the selection at the growing edge of the colony is for fastest growing individuals; while the microenvironment at the centre of the colony comprises of environmental (solvent) and nutritional stresses. Continuous propagation of MA lines, while selecting cells from the edge or centre of a colony are likely to yield different phenomenology. We explicitly test this hypothesis.

Thirty-two lines were propagated with cells from the edge of a colony were used to transfer from one plate to another. Similarly, thirty-two lines were propagated where cells from the centre of a colony were used to transfer from one plate to the next. After a 100 transfers, the mean growth rates of the three large lines (l1-l32), (c1-c32), and (e1-e32) were compared. The mean growth rate of the (e1-e32) lines were the largest and that of the c1-c32 lines were the least, and that of the lines l1-l32 between the two (**Figure 8A**). This demonstrates the action of selection in the growing colony. Moreover, when the cells from the lines were subjected to a heat shock for one hour (see methods for more details), the survival rate for the cells from centre lines (c1-c32) was found to be higher than that from the edge lines (e1-e32) (**Figure 8B**). As a result, the optical density in the c1-c32 lines, after 4 hours of growth post-shock, was higher than that in the edge lines (e1-e32) (**Figure 8C**).

**Figure 8.**
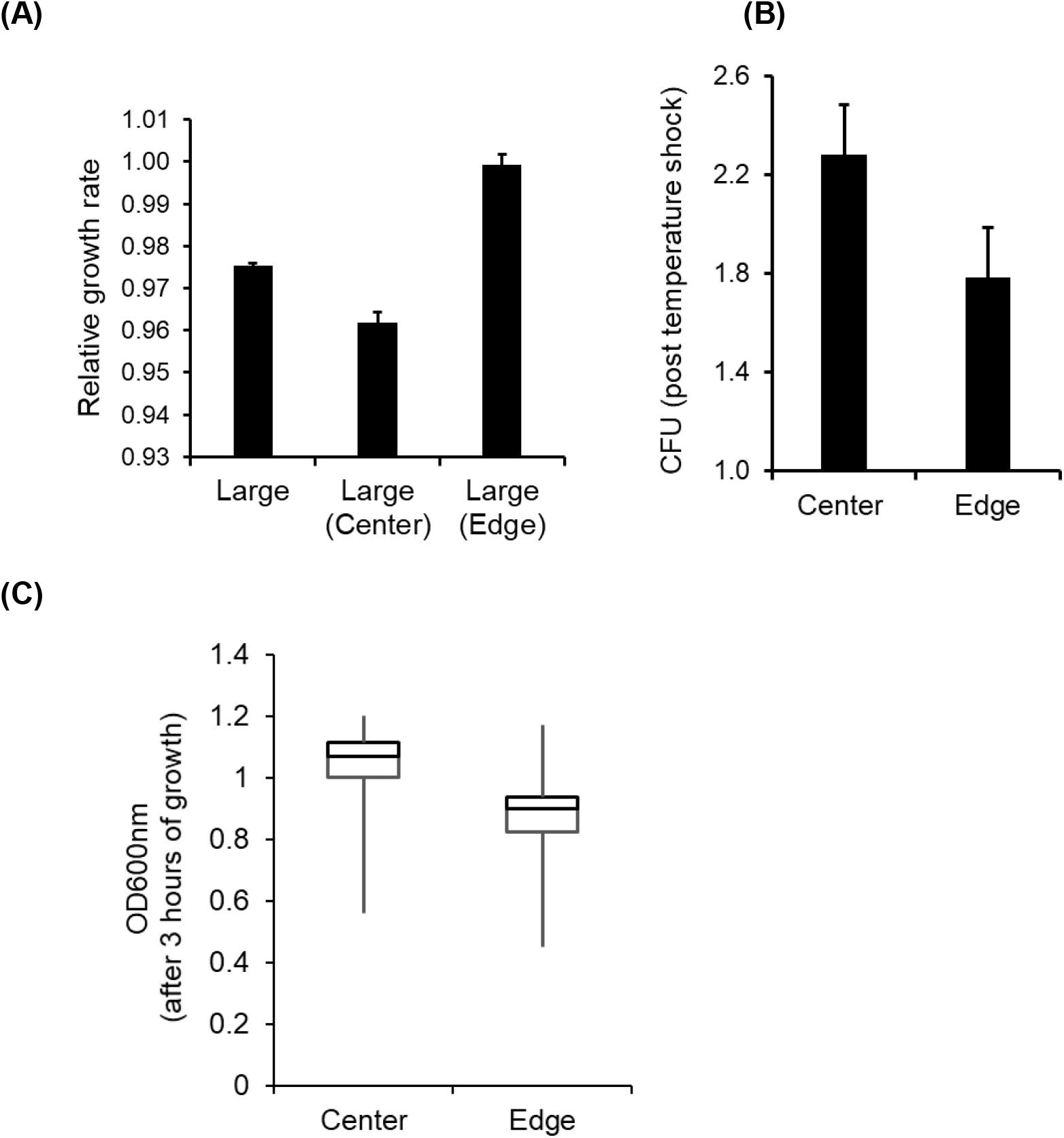
**(A)** Relative growth rate of the lines Large, Large-edge, and Large-center, with respect to the growth rate of the ancestor. The mean of three independent experiments of all 32 lines, and the standard error is represented. **(B)** Mean CFU count in the centre lines (c1-c32) is statistically significantly higher than that in the edge lines (e1-e32), when spotted for single colonies after heat shock at 50 deg C for one hour (t-test, p-value < 0.2). **(C)** Box-plot of ODs of the 32 centre and edge lines after 4 hours of growth at 37 deg C, after the heat shock. The mean OD of the centre lines is greater than that of the edge lines (t-test, p-value < 0.0001).

## Discussion

The idea that selection acts during an MA experiment conducted with bacteria/yeast was previously suggested by (Daniel L. Halligan, 2009). In a recent study, it was demonstrated with *Pseudomonas* that large beneficial mutations can impact fitness trajectories in an MA experiment (Heilbron et al., 2014). It was also recently demonstrated that both, the spectrum of mutations and the mutation rate in yeast is a function of the environment in which the organism is propagated (Liu & Zhang, 2019). However, how selection dictates this dynamics has not been previously addressed.

In this work, we address the question of how does selection impact the results from an MA experiment, and how does this change our analysis from the observations from these experiments. We demonstrate that, during a mutation accumulation experiment with bacteria or yeast, selection is able to act to a sufficient degree to significantly bias the nature of the mutations that fix in the population (Daniel L. Halligan, 2009). The bottleneck of population size one between two transfers does not eliminate the influence of selection during the growth phase of the colony. We show that the assumptions that selection in the colony is minimal since most cell divisions take place in the last generation (Lee et al., 2012) or that repeated bottlenecks in an MA experiment makes selection redundant (Katju & Bergthorsson, 2019), are not valid during the growth phase of the colony. In fact, selection acts quite strongly in an MA experiment, and as a result, on an average, beneficial mutations are more than twice as much represented as would be expected in the absence of selection. This is perhaps not surprising. In the context of growth of a colony, starting from a single colony to a population size of ~10^7^ constitutes ~25 generations. The effective population size of the population in this phase is ~10 (Zeyl et al., 2001).

As shown in our results, the estimates of mutation rates, DFEs, trends of mutation rates obtained from MA experiments are biased by selection. The manifested mutations are a result of selection and chance, and this process is dictated by several parameters, which feed into our model. These factors include mutation rate, fraction of mutations that are beneficial, DFE of beneficial and deleterious. The problem of deciphering mutation rate (or other parameters) from phenotypic or genotypic data from an MA experiment then becomes an inverse problem, such as one attempted previously in the context of DFE of beneficial mutations (Hegreness et al., 2006).

Moreover, a colony is a highly heterogeneous environment, which leads to different selection pressures and phenotypic manifestations of the participating individuals (Varahan et al., 2019, Cap et al., 2012, van Gestel & Nowak, 2016, van Boxtel et al., 2017). The fitness trajectories of an MA experiment, when propagated from the edge of a colony would be quite different from an MA experiment where propagation is done from the center of the colony. For instance, in yeast and bacteria, local environments within a colony are known to elicit different physiological responses (Carmona-Gutierrez et al., 2010, Jeanson et al., 2015, Mikkelsen et al., 2007). Similarly, the size of the colony at which propagation is done is also an important parameter. CFU increase in a growing colony is initially exponential with time, then linear, and eventually decreases to zero (data not shown). This implies that, in addition to different selection pressure in different regions of a colony, the relative strengths of drift and selection are also changing with time in a growing colony.

## Acknowledgements

AM is supported by the Council of Scientific and Industrial Research (CSIR), Government of India, as a Senior Research Fellow (09/087(0873)/2017-EMR-I). This research was supported in part by the International Centre for Theoretical Sciences (ICTS) during a visit (AM and SS) for participating in the program - Fourth Bangalore School on Population Genetics and Evolution (ICTS/popgen2020/01).

The authors thank Paike Jayadeva Bhat for discussions.

## Author contributions

AM: conceived the study, performed MA experiments, designed the simulations, analysed data, and wrote manuscript.

SK: performed simulations and analysed the data.

SS: conceived the study, analysed data, and wrote manuscript.

## References

Andersson, D. I. & Hughes, D. 1996. Muller’s ratchet decreases fitness of a DNA-based microbe. Proc Natl Acad Sci U S A 93: 906–7.

Ann-Marie Waldvogel, M. P. 2020. Temperature-dependence of spontaneous mutation rates. bioRxiv.

Baer, C. F., Shaw, F., Steding, C., Baumgartner, M., Hawkins, A., Houppert, A., Mason, N., Reed, M., Simonelic, K., Woodard, W. & Lynch, M. 2005. Comparative evolutionary genetics of spontaneous mutations affecting fitness in rhabditid nematodes. Proc Natl Acad Sci U S A 102: 5785–90.

Bondel, K. B., Kraemer, S. A., Samuels, T., McClean, D., Lachapelle, J., Ness, R. W., Colegrave, N. & Keightley, P. D. 2019. Inferring the distribution of fitness effects of spontaneous mutations in Chlamydomonas reinhardtii. PLoS Biol 17: e3000192.

Bosshard, L., Dupanloup, I., Tenaillon, O., Bruggmann, R., Ackermann, M., Peischl, S. & Excoffier, L. 2017. Accumulation of Deleterious Mutations During Bacterial Range Expansions. Genetics 207: 669–684.

Brajesh, R. G., Dutta, D. & Saini, S. 2019. Distribution of fitness effects of mutations obtained from a simple genetic regulatory network model. Sci Rep 9: 9842.

Caballero, A. & Keightley, P. D. 1998. Inferences on genome-wide deleterious mutation rates in inbred populations of Drosophila and mice. Genetica 102-103: 229–39.

Cap, M., Stepanek, L., Harant, K., Vachova, L. & Palkova, Z. 2012. Cell Differentiation within a Yeast Colony: Metabolic and Regulatory Parallels with a Tumor-Affected Organism. Molecular Cell 46: 436–448.

Cap, M., Vachova, L. & Palkova, Z. 2009. Yeast colony survival depends on metabolic adaptation and cell differentiation rather than on stress defense. J Biol Chem 284: 32572–81.

Cap, M., Vachova, L. & Palkova, Z. 2010. How to survive within a yeast colony?: Change metabolism or cope with stress? Commun Integr Biol 3: 198–200.

Carmona-Gutierrez, D., Eisenberg, T., Buttner, S., Meisinger, C., Kroemer, G. & Madeo, F. 2010. Apoptosis in yeast: triggers, pathways, subroutines. Cell Death and Differentiation 17: 763–773.

Daniel L. Halligan, P. D. K. 2009. Spontaneous Mutation Accumulation Studies in Evolutionary Genetics. Annual Review of Ecology, Evolution, and Systematics 40: 151–172.

Denver, D. R., Wilhelm, L. J., Howe, D. K., Gafner, K., Dolan, P. C. & Baer, C. F. 2012. Variation in base-substitution mutation in experimental and natural lineages of Caenorhabditis nematodes. Genome Biol Evol 4: 513–22.

Dickinson, W. J. 2008. Synergistic fitness interactions and a high frequency of beneficial changes among mutations accumulated under relaxed selection in Saccharomyces cerevisiae. Genetics 178: 1571–8.

Dillon, M. M. & Cooper, V. S. 2016. The Fitness Effects of Spontaneous Mutations Nearly Unseen by Selection in a Bacterium with Multiple Chromosomes. Genetics 204: 1225–1238.

Dillon, M. M., Sung, W., Lynch, M. & Cooper, V. S. 2018. Periodic Variation of Mutation Rates in Bacterial Genomes Associated with Replication Timing. mBio 9.

Draghi, J. A., Parsons, T. L., Wagner, G. P. & Plotkin, J. B. 2010. Mutational robustness can facilitate adaptation. Nature 463: 353–5.

Elena, S. F., Ekunwe, L., Hajela, N., Oden, S. A. & Lenski, R. E. 1998. Distribution of fitness effects caused by random insertion mutations in Escherichia coli. Genetica 102-103: 349–58.

Estes, S., Phillips, P. C., Denver, D. R., Thomas, W. K. & Lynch, M. 2004. Mutation accumulation in populations of varying size: the distribution of mutational effects for fitness correlates in Caenorhabditis elegans. Genetics 166: 1269–79.

Gautam Reddy, M. M. D. 2020. Global epistasis emerges from a generic model of a complex trait. bioRxiv.

Good, B. H. & Desai, M. M. 2015. The impact of macroscopic epistasis on long-term evolutionary dynamics. Genetics 199: 177–90.

Haag-Liautard, C., Dorris, M., Maside, X., Macaskill, S., Halligan, D. L., Houle, D., Charlesworth, B. & Keightley, P. D. 2007. Direct estimation of per nucleotide and genomic deleterious mutation rates in Drosophila. Nature 445: 82–5.

Hall, D. W., Fox, S., Kuzdzal-Fick, J. J., Strassmann, J. E. & Queller, D. C. 2013. The rate and effects of spontaneous mutation on fitness traits in the social amoeba, Dictyostelium discoideum. G3 (Bethesda) 3: 1115–27.

Hegreness, M., Shoresh, N., Hartl, D. & Kishony, R. 2006. An equivalence principle for the incorporation of favorable mutations in asexual populations. Science 311: 1615–7.

Heilbron, K., Toll-Riera, M., Kojadinovic, M. & MacLean, R. C. 2014. Fitness is strongly influenced by rare mutations of large effect in a microbial mutation accumulation experiment. Genetics 197: 981–90.

Jeanson, S., Floury, J., Gagnaire, V., Lortal, S. & Thierry, A. 2015. Bacterial Colonies in Solid Media and Foods: A Review on Their Growth and Interactions with the Micro-Environment. Frontiers in Microbiology 6.

Katju, V. & Bergthorsson, U. 2019. Old Trade, New Tricks: Insights into the Spontaneous Mutation Process from the Partnering of Classical Mutation Accumulation Experiments with High-Throughput Genomic Approaches. Genome Biol Evol 11: 136–165.

Keightley, P. D. & Caballero, A. 1997. Genomic mutation rates for lifetime reproductive output and lifespan in Caenorhabditis elegans. Proc Natl Acad Sci U S A 94: 3823–7.

Keightley, P. D., Trivedi, U., Thomson, M., Oliver, F., Kumar, S. & Blaxter, M. L. 2009. Analysis of the genome sequences of three Drosophila melanogaster spontaneous mutation accumulation lines. Genome Res 19: 1195–201.

Kibota, T. T. & Lynch, M. 1996. Estimate of the genomic mutation rate deleterious to overall fitness in E. coli. Nature 381: 694–6.

Kraemer, S. A., Bondel, K. B., Ness, R. W., Keightley, P. D. & Colegrave, N. 2017. Fitness change in relation to mutation number in spontaneous mutation accumulation lines of Chlamydomonas reinhardtii. Evolution 71: 2918–2929.

Krasovec, M., Eyre-Walker, A., Sanchez-Ferandin, S. & Piganeau, G. 2017. Spontaneous Mutation Rate in the Smallest Photosynthetic Eukaryotes. Mol Biol Evol 34: 1770–1779.

Kryazhimskiy, S., Rice, D. P., Jerison, E. R. & Desai, M. M. 2014. Microbial evolution. Global epistasis makes adaptation predictable despite sequence-level stochasticity. Science 344: 1519–1522.

Kuroki, S., Ohta, A., Katoh, O., Sueko, N., Yamada, H. & Yamaguchi, M. 1993. Successful treatment of autoimmune thrombocytopenia and rapidly progressive pneumonitis with recurrent pneumothoraces in a patient with rheumatoid arthritis. Br J Rheumatol 32: 855–6.

Lee, H., Popodi, E., Tang, H. & Foster, P. L. 2012. Rate and molecular spectrum of spontaneous mutations in the bacterium Escherichia coli as determined by whole-genome sequencing. Proc Natl Acad Sci U S A 109: E2774–83.

Liu, H. & Zhang, J. 2019. Yeast Spontaneous Mutation Rate and Spectrum Vary with Environment. Curr Biol 29: 1584–1591 e3.

Lynch, M., Sung, W., Morris, K., Coffey, N., Landry, C. R., Dopman, E. B., Dickinson, W. J., Okamoto, K., Kulkarni, S., Hartl, D. L. & Thomas, W. K. 2008. A genome-wide view of the spectrum of spontaneous mutations in yeast. Proc Natl Acad Sci U S A 105: 9272–7.

Mikkelsen, H., Duck, Z., Lilley, K. S. & Welch, M. 2007. Interrelationships between colonies, biofilms, and planktonic cells of Pseudomonas aeruginosa. Journal of Bacteriology 189: 2411–2416.

Mrudula Sane, G. D. D., Bhoomika A Bhat, Lindi M Wahl, Deepa Agashe 2020. Shifts in mutation spectra enhance access to beneficial mutations. bioRxiv.

Mukai, T. 1964. The Genetic Structure of Natural Populations of Drosophila Melanogaster. I. Spontaneous Mutation Rate of Polygenes Controlling Viability. Genetics 50: 1–19.

Mukai, T., Chigusa, S. I., Mettler, L. E. & Crow, J. F. 1972. Mutation rate and dominance of genes affecting viability in Drosophila melanogaster. Genetics 72: 335–55.

Neher, R. A. 2013. Genetic Draft, Selective Interference, and Population Genetics of Rapid Adaptation. Annual Review of Ecology, Evolution, and Systematics 44: 195–215.

Ness, R. W., Morgan, A. D., Colegrave, N. & Keightley, P. D. 2012. Estimate of the spontaneous mutation rate in Chlamydomonas reinhardtii. Genetics 192: 1447–54.

Orr, H. A. 2010. The population genetics of beneficial mutations. Philos Trans R Soc Lond B Biol Sci 365: 1195–201.

Ossowski, S., Schneeberger, K., Lucas-Lledo, J. I., Warthmann, N., Clark, R. M., Shaw, R. G., Weigel, D. & Lynch, M. 2010. The rate and molecular spectrum of spontaneous mutations in Arabidopsis thaliana. Science 327: 92–4.

Perfeito, L., Fernandes, L., Mota, C. & Gordo, I. 2007. Adaptive mutations in bacteria: high rate and small effects. Science 317: 813–5.

Pugatch, R. 2015. Greedy scheduling of cellular self-replication leads to optimal doubling times with a log-Frechet distribution. Proc Natl Acad Sci U S A 112: 2611–6.

Rutter, M. T., Roles, A., Conner, J. K., Shaw, R. G., Shaw, F. H., Schneeberger, K., Ossowski, S., Weigel, D. & Fenster, C. B. 2012. Fitness of Arabidopsis thaliana mutation accumulation lines whose spontaneous mutations are known. Evolution 66: 2335–9.

Schrider, D. R., Houle, D., Lynch, M. & Hahn, M. W. 2013. Rates and genomic consequences of spontaneous mutational events in Drosophila melanogaster. Genetics 194: 937–54.

van Boxtel, C., van Heerden, J. H., Nordholt, N., Schmidt, P. & Bruggeman, F. J. 2017. Taking chances and making mistakes: non-genetic phenotypic heterogeneity and its consequences for surviving in dynamic environments. Journal of the Royal Society Interface 14.

van Gestel, J. & Nowak, M. A. 2016. Phenotypic Heterogeneity and the Evolution of Bacterial Life Cycles. Plos Computational Biology 12.

Varahan, S., Walvekar, A., Sinha, V., Krishna, S. & Laxman, S. 2019. Metabolic constraints drive self-organization of specialized cell groups. Elife 8.

Wagner, A. 2005. Robustness, evolvability, and neutrality. FEBS Lett 579: 1772–8.

Wagner, A. 2008. Neutralism and selectionism: a network-based reconciliation. Nat Rev Genet 9: 965–74.

Wagner, A. 2012. The role of robustness in phenotypic adaptation and innovation. Proc Biol Sci 279: 1249–58.

Wang, P., Robert, L., Pelletier, J., Dang, W. L., Taddei, F., Wright, A. & Jun, S. 2010. Robust growth of Escherichia coli. Curr Biol 20: 1099–103.

Wrenbeck, E. E., Azouz, L. R. & Whitehead, T. A. 2017. Single-mutation fitness landscapes for an enzyme on multiple substrates reveal specificity is globally encoded. Nat Commun 8: 15695.

Zeyl, C. & DeVisser, J. A. 2001. Estimates of the rate and distribution of fitness effects of spontaneous mutation in Saccharomyces cerevisiae. Genetics 157: 53–61.

Zeyl, C., Mizesko, M. & de Visser, J. A. 2001. Mutational meltdown in laboratory yeast populations. Evolution 55: 909–17.

